# Reconstructing cancer karyotypes from short read data: the half full and half empty glass

**DOI:** 10.1101/152447

**Authors:** Rami Eitan, Ron Shamir

**Affiliations:** Blavatnik School of Computer Science, Tel Aviv University, Tel Aviv, 69978 Israel

**Keywords:** cancer, karyotypes, genome rearrangements, structural and numerical variations, deep sequencing, reconstruction, graph theory, integer linear programming

## Abstract

**Background:** During cancer progression genomes undergo point mutations as well as larger segmental changes. The latter include, among others, segmental deletions duplications, translocations and inversions. The result is a highly complex, patient-specific cancer karyotype. Using high-throughput technologies of deep sequencing and microarrays it is possible to interrogate a cancer genome and produce chromosomal copy number profiles and a list of breakpoints (“jumps”) relative to the normal genome. This information is very detailed but local, and does not give the overall picture of the cancer genome. One of the basic challenges in cancer genome research is to use such information to infer the cancer karyotype.

We present here an algorithmic approach, based on graph theory and integer linear programming, that receives segmental copy number and breakpoint data as input and produces a cancer karyotype that is most concordant with them. We used simulations to evaluate the utility of our approach, and applied it to real data.

**Results:** By using a simulation model, we were able to estimate the correctness and robustness of the algorithm in a spectrum of scenarios. Under our base scenario, designed according to observations in real data, the algorithm correctly inferred 69% of the karyotypes. However, when using less stringent correctness metrics that account for incomplete and noisy data, 87% of the reconstructed karyotypes were correct. Furthermore, in scenarios where the data were very clean and complete, accuracy rose to 90%-100%. Some examples of analysis of real data, and the karyotypes reconstructed by our algorithm, are also presented.

**Conclusion:** While reconstruction of complete, perfect karyotype based on short read data is very hard, a large portion of the reconstruction will still be correct and can provide useful information.

## Background

The current understanding of cancer suggests that it is a disease driven by somatic mutations that accumulate in the genome, within a certain tissue, during the lifetime of an individual. These mutations vary in size and effect. They can be small, e.g., single nucleotide mutations, or large structural variations caused by rearrangements such as deletions, inversions, tandem duplications and chromosomal translocations, or duplication and losses of entire chromosomes ^1^. Over time these rearrangements accumulate and result in genomes less and less similar to the germline genome.

Cancer genomes are often described in the form of karyotypes. A *karyotype* is a high level description of the genome as a set of chromosomes and the number of copies of each. Normal karyotypes have two copies of each chromosome 1 to 22 and the sex chromosomes. In contrast, in cancer karyotypes some chromosomes may contain fragments originating from several normal chromosomes.

### Types of aberration events

Most segmental changes that happen during the progression of the disease can be categorized as deletion, tandem duplication, inversion, translocation, and deletion and duplication of entire chromosomes.

A *deletion* is characterized by a missing segment of a chromosome, a *tandem duplication* happens when part of the chromosome is duplicated and thus two copies of a segment appear where normally there would only be one. An *inversion* occurs when a segment of a chromosome is reversed relative to its original orientation (Figure 0).

**Figure 0:**
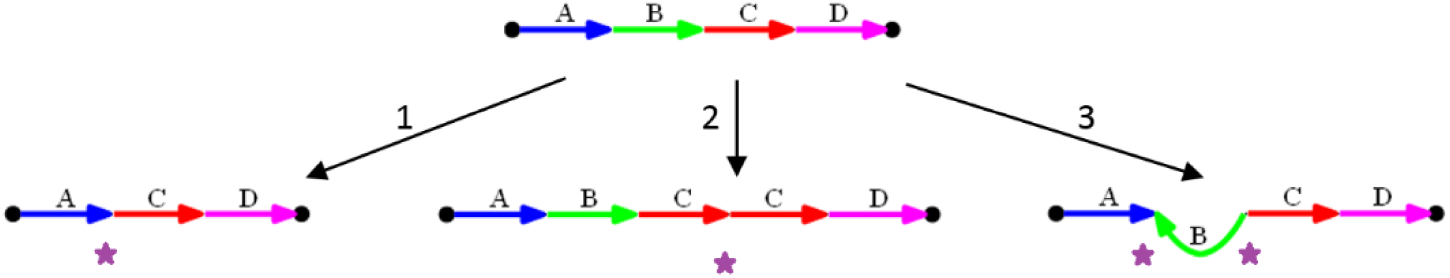
Basic types of rearrangements. (1) Deletion: segment B of the normal chromosome is deleted. (2) Tandem duplication: segment C duplicates and repeats. (3) Inversion: segment B is inverted. Stars indicate breakpoints.

A *translocation* happens when two different chromosomes “switch” end segments. Schematically, a translocation on two chromosomes (A,B) and (C,D) produces the chromosomes (A,D) and (C,B). A *whole chromosomal duplication* (*deletion*) adds (removes) a copy of a complete chromosome.

### Breakpoints

The molecular mechanisms that cause somatic genome rearrangements are still the focus of investigation. The main paradigm is that a genome rearrangement occurs when one or more chromosomes break and a following joining event reassembles the fragments in a different order. A *breakpoint* is defined as a genomic location where the normal DNA sequence is interrupted and two non-adjacent sequence segments appear consecutively due to a joining event. A breakpoint can be considered as the most basic unit of rearrangement. The stars in Figure 0 indicate breakpoints.

### Models of genomic distance

Modeling the somatic evolution of cancer holds great value for understanding the disease process. In 1995 Hannenhalli and Pevzner proposed a method to calculate the genomic distance between two species based on the minimal number of reversals (or reversals and translocations, in the multi-chromosome case) required to transform the genome of one species to another ^2,3^.

Braga et al. proposed another distance metric between genomes. They developed a method that calculates the distance between two genomes based on *Double-Cut and Join* (*DCJ*) operations and *indels* (Insertions and deletions), and utilized it to show evidence for deletion clusters in six species of Rickettsia ^4^. Feijão et al. defined another metric based on *Single-cut and join* (*SCJ*) operations, and by using it they were able to recover between 60 and 90 percent of the topology of a phylogenic tree with 200 different genomes and with as many as 3000 genes ^5,6^. Zeira and Shamir defined a generalized model, called *SCJD*, allowing the operations of cut, join and whole chromosome duplication. They developed a linear time algorithm for computing the shortest sequence of operations transforming one linear genome with one copy per gene into another with two copies per gene ^7^.

Ozery-Flato and Shamir introduced the *elementary distance* between two karyotypes, defined as the least number of elementary operations – breakage, fusion, duplication and deletion – transforming one into the other. They suggested a polynomial time 3-approximation algorithm to find the shortest elementary distance between two karyotypes. Applying the algorithm on some 58,000 karyotypes taken from the Mitelman database ^8^, 99.9% of the resulting solutions matched the lower (optimal) bound ^9^.

### Detecting chromosomal aberrations

#### Paired end reads

One of the main ways for inferring breakpoints in the genome, detecting structural variants and identifying rearrangements is using *paired end reads* produced by deep sequencing ^10–13^. Paired end reads are generated by fragmenting the genomic DNA into short segments, followed by sequencing both ends of (some of) the segments (Figure 1). Typical lengths are ~350 bp per fragment (also called *insert*) and ~100 bp per read (end). The unsequenced segment of the insert is called the *gap* (length ~150 bp in the example above). The two ends of each read are then aligned back to a reference normal genome (in the case of cancer – the genome of a healthy cell from the same patient). The approximate length of the insert and the relative orientation of its ends is known in advance. We expect the two ends of a fragment to be aligned to the reference genome at roughly that distance and with the correct relative orientation. An alignment is called a *concordant* if it meets those conditions, and *discordant* otherwise.

**Figure 1:**
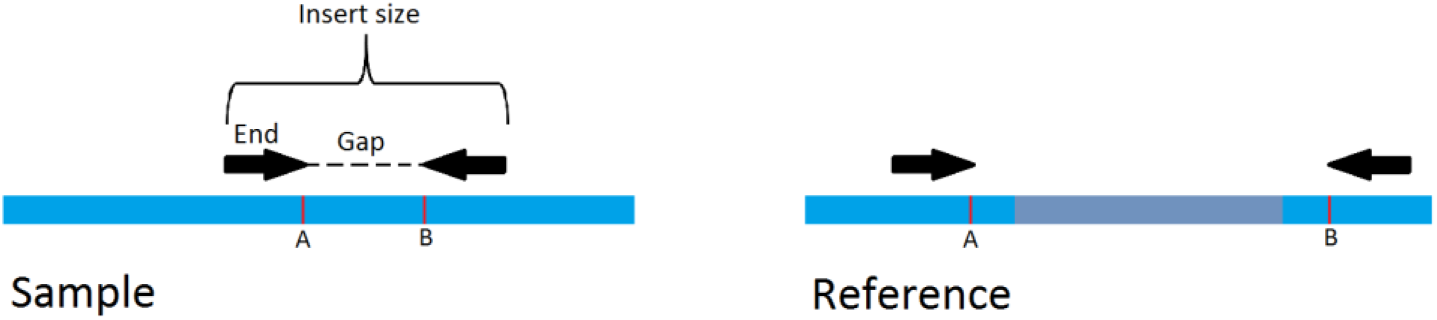
The paired ends read alignment signature of a deletion rearrangements. The grey area on the reference genome between points A and B was deleted in the sample genome. Any read whose gap falls between A and B on the sample genome will have its ends aligned to locations that are far apart on the same chromosome, indicating a deletion. Other rearrangements leave unique signatures in a similar manner.

Discordant reads suggest a breakpoint in the genome. A read taken from that spot will have its two ends aligned to locations on the reference genome where those positions originally lie. The type of discordance suggests the rearrangement event that occurred (See ^14^).

#### Detecting structural variations

A first step in analysis of paired end reads is their mapping to the reference genome. A variety of computational approaches were developed for inferring the structural variations from the discordant reads and produce a set of rearrangement events ^15–20^. Other methods such as PREGO ^21^ take into account the concordant reads as well. BreaKmer ^14^ uses the misaligned reads together with the aligned concordant and discordant reads to predict rearrangements using k-mer statistics. CouGaR ^22^ is a method for identifying large-scale complex genomic rearrangements using both depth of coverage and discordant paired-ends mapping. SV-Bay ^23^ applies a Bayesian approach to data of mapped paired-ends reads to infer breakpoint locations and copy number variations and predict structural variations in a cancer genome. Recently, a new algorithm, Weaver ^24^, was proposed to estimate both the allelic copy number and inter-connectivity of SV’s using a probabilistic graph model. Expanding on Weaver, Rajaraman et al. ^25^ used a graph model and an ILP formulation to further predict SV phasing and the interconnectivity of unphased SV’s with high specificity. Other algorithmic approaches infer rearrangements that are less simple and have more complex signatures ^14,26,27^.

Some methods seek to achieve higher accuracy by aggregating results from several different tools. MetaSV ^28^ offers an improvement of accuracy and precision in detecting different kinds of structural variants. By effectively merging the results from multiple tools, they were able to reach F1-scores (harmonic mean of sensitivity and precision) of 96.2% for deletions and 84.7% for insertions. SomaticSeq ^29^ detects single nucleotide variants (SNVs) and small insertions and deletions (indels), using machine learning algorithms to incorporate the results from five somatic mutation callers. The authors report an F1 score of 90%.

### Copy number variations

Duplications and deletions change the copy number (CN) of different segments of the DNA sequence, i.e. the number of times a segment is present in the karyotype. A normal (human) cell line has 22 diploid chromosomes (ignoring the sex chromosomes XX or XY) and so the CN of the entire karyotype is 2. A gain or a loss of an entire chromosome will decrease or increase the CN of that chromosome, respectively. A fraction of a chromosome can also be deleted or duplicated. The resulting segment or chromosome is said to have undergone a *copy number variation* (CNV).

Large CNVs can be detected by traditional methods like Fluorescence in-situ Hybridization (FISH) ^30^. Higher resolution detection of CNVs can be achieved by Array Comparative Genomic Hybridization (aCGH) ^31^. With the advent of next-generation sequencing (NGS), several methods have been developed to infer CNV’s using DNA sequences ^21,32,33^. NGS based methods have the potential to greatly increase the resolution of CNV analysis, but they present many computational challenges and different methods may still vary widely in the results they produce on the same DNA sequence ^34^.

### Graph models for rearrangements

Graph theory has been highly instrumental in the area of genomic rearrangements. For example, de Bruijn graphs are used for genome assembly problems ^35^, and breakpoint graphs are used in reconstructing rearranged genomes across species ^2,36^. More recently, similar methods were adapted for cancer genomes ^9,37^. The breakpoint graph, introduced by Pevzner and Bafna in 1993 to represent the relation between two permutations of the same set of elements ^38^, remains today one of the key models in the study of genomic rearrangements. Greenman et al. expanded on the breakpoint graph and introduced a construction that is essentially equivalent called the *allelic graph* and its counterpart the *somatic graph* ^39^.

Oesper et al. proposed a construction that expands on the breakpoint graph, called interval adjacency graph ^21^. The interval adjacency graph is constructed directly from CN and breakpoint data. The discordant reads are used to infer breakpoint locations on the DNA sequence and partition it to intervals accordingly. A full description of the graph appears in Section 0.

Using the interval adjacency graph it is possible to infer rearranged sequences that agree with the data. Oesper et al. showed that an Eulerian path on the graph alternating between interval edges and reference / variant edges corresponds to a rearranged sequence of the chromosome. They developed an algorithm called *PREGO* to determine the most likely sequence of a rearranged karyotype. Using simulations they showed their algorithm can deduce the correct multiplicity of more than 80% of the variant edges, even with high noise and when the sample is heterogeneous. Furthermore, they applied PREGO to five ovarian cancer genomes and were able to identify numerous rearrangements and structural variants, some of which were consistent with known mechanisms. PREGO combines CN and adjacency information from paired end reads to infer multiplicity of different segments in the cancer genome. However except in simple cases, the underlying karyotype cannot be uniquely resolved, as many reconstructions will be consistent with the data.

## Methods

We propose here a novel method that receives as input discordant paired-end reads and genomic CNs obtained from sequencing a cancer genome, and reconstructs a karyotype that is in most agreement with the input. The outline of our approach is as follows. We use the two data types together to construct a *bridge graph*, akin to the adjacency graph proposed by Oesper et al. ^21^. An integer linear programming (ILP) optimization problem is formulated and then solved on the graph. The solution is a valid karyotype of the rearranged genome that is most concordant with the observed data. We also present the solution graphically.

### The adjacency and bridge graphs

In our problem setup there is a *normal* (*or reference*) *genome*, whose contents is known, and an unknown *target genome* that should be reconstructed. A *breakpoint* is a point along the reference genome involved in a structural change event in the target genome.

Let *C* be the set of chromosomes in the reference karyotype. The breakpoints partition each chromosome *c* ∊ *C* into a set of *k^c^* intervals 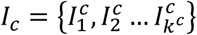, such that each 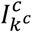 is an interval between consecutive breakpoints, or between a breakpoint and a chromosome end. The intervals are numbered in increasing order along *c*, so that *c* is equal to the concatenation of the intervals 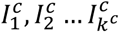. We call the start and end points of interval *I* the *tail* and *head* of *I* and denote them by *t_I_* and *h_I_* respectively. Hence, *I =* [*t_I_*,*h_I_*], and −*I =* [*h_I_*,*t_I_*] is the interval *I* reversed. An *extremity* is a tail or a head of an interval. The set of all intervals 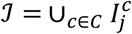constitutes the set of the basic building blocks of the reference and target genomes. The length of interval *I_j_* (in bases) is denoted by *l_j_*, and 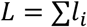 is the total length of all intervals.

The target genome can be represented by a set of chromosomes, where each chromosome is a sequence of intervals, some possibly reversed (Figure 3). A *bridge* is a pair of extremities that are not adjacent on the reference genome but are adjacent in the target genome. Bridges can be detected based on the paired-end read data of the target genome (Figure). The *support level* of bridge *b_i_* is the number of paired-end reads that support it, denoted *μ_i_*. The total support score for all bridges is denoted 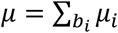.

**Figure 3:**
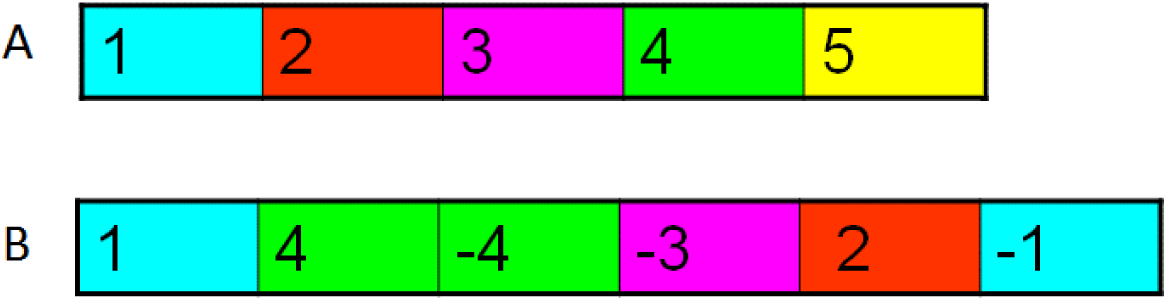
Reference and target genomes. A: reference (germline) chromosome segmented into intervals separated by breakpoints. B: The rearranged chromosome represented by the series of intervals 1,4,-4,-3,2,-1. Genome B contains the bridges {h_1_, t_4_}, {h_4_, h_4_}, {t_3_, t_2_} and {h_2_, h_4_}. Note that {t_4_, h_3_} is not a bridge.

Each interval *I_i_* ∊ *I* has a *CN N_i_* ≥ 0 indicating the number of times it appears in the target genome. The set of CNs of all intervals is called the *copy number profile* of the target. That profile can be derived from deep sequencing data or from array CGH data. In perfect data, *N_i_* is exactly the number of copies of the interval in the target genome. In practice, the CNs are real valued estimates based on mean coverage of each interval.

Let us first reiterate the definition of the *interval adjacency graph*, introduced in ^21^. The input is (1) the reference genome represented as a sequence of intervals for each chromosome. These intervals form the set *J =* {*I*_1_, …, *I_n_*}; interval *I_j_* has length *l_j_*. (2) The CN profile of the intervals: Interval *I_j_* has CN *N_j_*. (3) The set of bridges 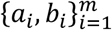 and the support *μ_i_* for each bridge. Each *a_i_* and *b_i_* is an extremity of an interval in J. We define a weighted undirected graph *G*(*V*, *E*, *w*) whose vertices are the interval extremities. For each interval *I_i_* = [*t_i_, h_i_*], the graph contains an *interval edge e_I_*(*t_i_*, *h_i_*) ∊ *E_I_* connecting its two extremities, of weight *N_i_*. For each two intervals *I_i_*, *I_i+_*_1_ that are adjacent on the reference genome, a *reference edge e_R_*(*h_i_, t_i+_*_1_) ∊ *E_R_* connects the head of *I_i_* to the tail of *I_i+_*_1_. Reference edges are unweighted. Each bridge is represented by a *bridge* (*or variant*) *edge e_V_*(*a_i_*, *b_j_*) ∊ *E_V_* connecting the two extremities *a_i_* and *b_j_*, with weight *μ_i_*. In total, the edge set of the graph is *E = E_I_* ∪*E_R_* ∪ *E_V_*. We denote by *S* ⊆*V* the set of vertices that represent *telomere nodes*, i. e. the nodes representing start and end points of each reference chromosome, hence 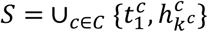 includes the heads of all starting intervals and the tails of all ending intervals in each chromosome’s partition.

A *bridge graph* is an interval adjacency graph with two minor changes: (1) bridge edges are assigned weights. The weight *w*(*e*) of the bridge *e*(*u*, *v*) is its *support score*, namely the number of paired end reads supporting that bridge. Hence, in a bridge graph both bridge and interval edges have weights. (2) We transform each undirected edge *e*(*u*,*v*) in the interval adjacency graph into two directed edges *e*_→_: *u* → *v*, *e*_←_: *v → u*. The original undirected edge is referred to as a *connection* to distinguish it from the directed edges, and *E = E*_→_ ∪ *E*_←_ is the set of edges in the graph. An example of a bridge graph is given in Figure 4.

**Figure 4:**
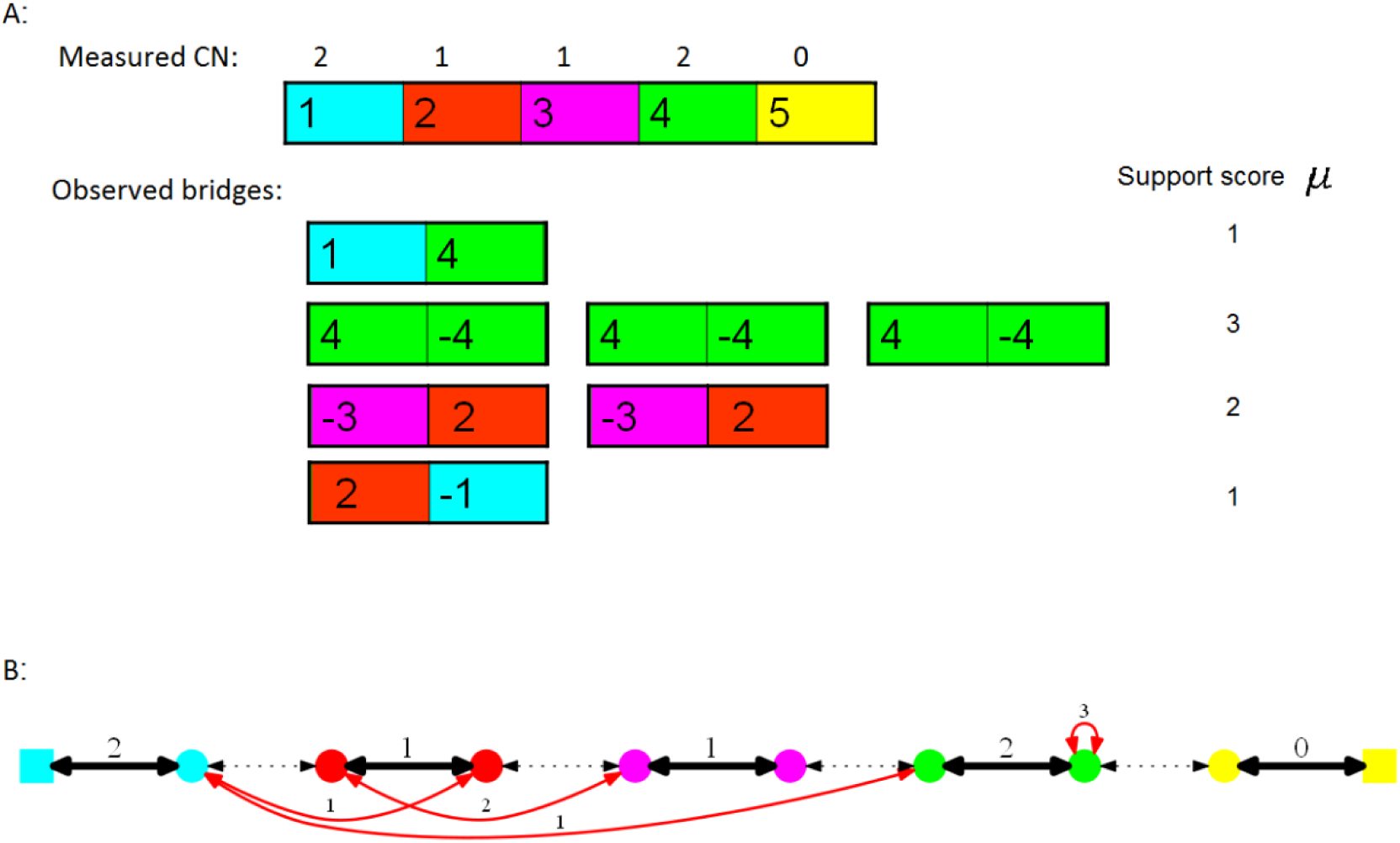
Bridge graph. The normal karyotype is the single chromosome (1,2,3,4,5). A: The measured CN and bridge data with the observed support score for each bridge. B: The corresponding bridge graph with weights for interval and bridge edges. All connections are composed of two antiparallel directed edges.

### Reconstructing the rearranged karyotype

Given the bridge graph *G*(*V*, *E*, *w*), we wish to find paths in *G* that correspond to rearranged chromosomes. Suppose first that the input data are complete and errorless. Recall that *S* ⊆ *V* is the set of vertices that represent *telomere nodes*, i.e. the nodes representing the start and end points of each chromosome. A *valid path p* is a path through *G* beginning and ending at *s*_1_, *s*_2_ ∊ *S* that alternately traverses interval and non-interval (i.e. reference/bridge) edges, and where the number of times each interval connection *e_i_* is traversed (in either direction), denoted *f_p_*(*e_i_*), is less than or equal to the CN of interval *i*, *N_i_*.

The requirement for an alternating path is because a traversal of an interval edge corresponds to traversing a segment from the reference genome, while a traversal of a reference/bridge edge is equivalent to a transition between segments. Therefore, such an alternating path represents a sequence of segments from the reference genome. Note that *f_p_*(*e_i_*) = *f_p_*(*e_i_*_→_) + *f_p_* (*e_i_*_←_) for every connection *e*. A set of such paths *P =* {*p*_1_, *p*_2_ … *p_n_*} where for each interval connection *e_i_*, Σ*_p_*_∊_*_P_ f_p_*(*e_i_*) = *N_i_* corresponds to a set of rearranged chromosomes, or a valid karyotype.

The restriction that the path alternates between interval and non-interval edges means that at each non-telomeric node *v* ∉ *S*, every traversal on an interval edge going into *v* must be followed by a traversal on a reference\bridge edge going out of *v*, and vice-versa. Telomeric nodes are excluded from this constraint as by definition they are the start or end of a path.

As detailed above, each connection between nodes *u,v* is composed of two antiparallel directed edges. For each node *v* ∊ *V* we denote *E*_*I*←_(*v*), *E*_*I*→_(*v*)*, E_R_*_←_(*v*), *E_R_*_→_(*v*)*, E_B_*_←_(*v*)*, E_B_*_→_(*v*) as the set of interval, reference and bridge edges that go in and out of *v* respectively. As above, we denote by *f_p_*(*e*) the number of times a connection *e* is traversed in path *p* and *f_P_*(*e*) = Σ*_p_*_∊_*_P_ f_p_*(*e*) is the total number of times a connection *e* is traversed in *P*. Additionally, for a set of connections *E*, *f_P_*(*E*) = Σ*_e_*_∊_*_E_ f_p_*(*e*) is the total number of times all connections in *E* are traversed in *P*. The constraints for a valid set of paths *P*, representing a rearranged karyotype, can be therefore formulated as:

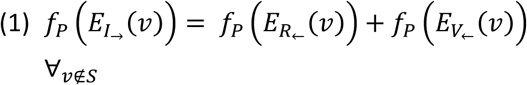

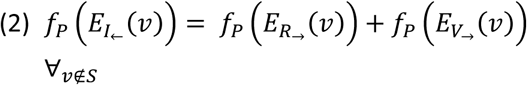

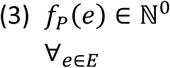

#### Scoring candidate solutions

Recall that the interval and bridge edges have weights, representing the measured CN of the intervals and the support score for the bridges, respectively. These values are in practice noisy. Given a bridge graph *G*(*V, E, w*) and a valid set of paths *P* representing a rearranged karyotype, we define a *discordance score* of *P*, denoted *d_G_*(*P*), which measures how much *P* is in agreement with the data in *G*, as follows:

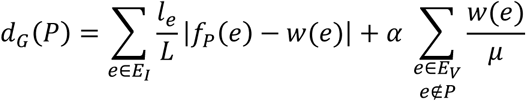

The first sum measures the disagreement of *P* with the CN profile. It is the sum over all interval edges *e* ∊ *E_I_* of the absolute difference between *f_P_*(*e*) and the input weight *w*(*e*), normalized by *l_e_*. We normalize the weights of the intervals by their lengths since longer genomic intervals are expected to have more accurate CN values, and hence should be penalized more for disagreement. Dividing by *L* guarantees that the range of the first sum is [0,1] if the absolute difference values are ≤ 1.

The second sum the disagreement of *P* with the bridge data. The more bridges *P* is utilizing, the more concordant it is with the bridge data. To reflect this, a penalty is given for each bridge edge *e* ∊ *E_V_* that is not used in *P*. The bigger the support score for a bridge is, the bigger the penalty if it is not used, and so the penalties are normalized by *w*(*e*). Dividing by 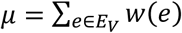 guarantees that the range of the second sum is [0,1]. To avoid summing over *e* ∉ *P*, we can rewrite the second term as 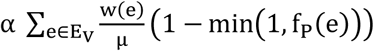.

The parameter *α* determines the relative weight the algorithm gives to paired-end reads data, i. e. how much it tries to utilize bridge edges in the solution. Using the algorithm on real tumor data, we set *α* = 0.5.

#### The ILP formulation

We wish to find a rearranged karyotype that is most consistent with the data, i.e., it corresponds to a valid set of paths and has smallest possible discordance score. This problem can be formulated as an ILP on the bridge graph *G*(*V, E*, *w*), as we now show.

For each connection *e_i_* ∊ *E* we define two variables *x_i→_*, *x_i_*_←_. The variables represent the number of times each edge is traversed in a path, and so *f_p_*(*e_i_*) *= x_i_*_→_ *+ x_i_*_←_. Each variable is noted *x^I^, x^B^* or *x^R^* for interval, bridge or reference edges respectively. Using these variables we can formulate the problem as follows.

**Minimize:**

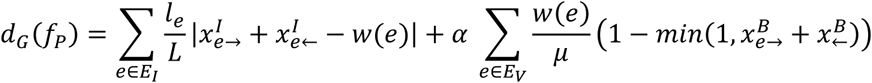

**Subject to:**

1. 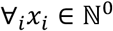
2. 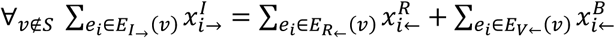
3. 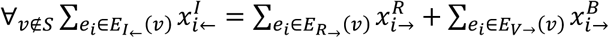

Constraint set (1) guarantees an integral non-negative solution. Constraints (2) and (3) are the valid path constraints. Note that telomeric nodes in *S* are not constrained.

### Tools

The core of the algorithm was implemented in java using the ILP solver package CPLEX, distributed by IBM ^40^ and was run on UNIX. The simulations module and the rest of the algorithm was implemented in python version 2.7 on Windows. The code is available in https://github.com/Shamir-Lab/Karyotype-reconstruction. A typical run of a single karyotype on a standard PC takes around 1 second.

## Results and discussion

### Simulations

To assess the performance of our algorithm, we simulated tumor karyotypes and applied the algorithm to them. To evaluate the quality of each reconstructed karyotype, it was compared to the correct karyotype, and summary statistics were computed. An overview of the simulation algorithm is as follows:

1. Start with a normal diploid karyotype *H* with *C* chromosomes
2. Perform *K* operations resulting in karyotype *T′*
3. Compute the exact (noiseless) CN profile and the bridges in *T′*
4. Add noise to the CN data and generate support values for the bridges

We start with a normal diploid karyotype *H* with a prescribed number of chromosomes. For simplicity, each chromosome is represented by a sequence of 300 atomic segments, which are its basic units. We perform a series of operations on the karyotype by applying deletions, inversions, tandem duplications and translocations. The types and the positions of the rearrangements are drawn uniformly at random. The span of operations that affect a single chromosomes (deletions, duplications and inversions) was limited to 30 atomic segments. This limit was set in order to avoid rapid erasure of large chromosomal segments by deletions. The total number of operations applied varies and determines the complexity of the resulting tumor karyotype *T*.

By comparing *H* and *T*, breakpoints are detected and each normal chromosome is partitioned into segments. Each segment has a CN (the number of occurrences of that segment in *T*). Each two consecutive segments in *T* that are not consecutive (and/or not in the same relative orientation) in *H* constitute a bridge. The clean (noiseless) data can thus be summarized as an integer-valued CN profile and the set of all bridges formed.

To simulate noisy scenarios, the CN profile and the bridge information is modified as follows. Normally distributed noise *x* is added to the CN of each segment independently, where *x ~ N*(*0*, *ϵ*). The support for each bridge (corresponding to the number of discordant reads supporting it) is drawn independently from an exponential distribution *Exp*(*λ*) (The exponential distribution was chosen based on empirical data with *λ* = 0.1866. See below). To simulate the possibility of bridges being completely missed, each bridge has probability *p* to completely be omitted from the final set of bridges.

In summary, the simulation program receives the following parameters (the default values appear in parentheses):

- *C* - The number of chromosomes (default: 5).
- *N* - The number of structural and numerical operations applied (default: 5).
- *ϵ* - The standard deviation of the noise in the CN profile data (default: 0.28)
- *p* – The probability to completely miss a bridge (default: 0.05).

In the *base scenario*, all parameters were at their default values. These parameters correspond to those computed on a tumor sample of medium complexity and a realistic level of noise (see Real tumor analysis below). Other scenarios were explored by changing the value of one of the parameters above while keeping the rest at their default levels.

#### Solution quality measures

We used five different measures for the level of correctness of a solution. Let *T* be the simulated (true) karyotype, let *T\** be the simulated noisy karyotype, and let *S* be the karyotype produced by the algorithm:

1. Is *S* equivalent to *T*? We say that *S* is *equivalent* to *T* if they have the same CN profile and both use the same bridges. Most equivalent karyotypes only differ in chromosomal orientation, and thus represent the same solution. We call such a solution *correct*.
2. Do *S* and *T* have the same CN profile? The CN of an interval is determined by many reads (or probes) and so is expected to be more robust than bridge information, determined by a few paired end reads. This criterion tests if *S* and *T* match in their CN profile. We call this criterion *Equal Copy Number* (ECN).
3. Does *S* have an equal or better score than *T?* When noise level is high, *T* and *T\** may differ substantially, and a solution closer to *T\** than to *T* does not indicate a failure of the algorithm but rather that the noise level is too high. Here the score is the ILP objective function value. We call this criterion *Equal or Better Score* (EBS).
4. Is *S* equivalent to *T* excluding missing bridges? *T\** may not include all the bridges found in *T*, and in that case *S* can never be equivalent to *T*. However, we consider *S* to be correct for all observed bridges if it has the correct CN profile for all segments that are unaffected by a missed bridge, and is using all the bridges from *T* that are included in *T*\*(Figure S2). We call this metric *Equivalent for Observed Bridges* (EOB).
5. What fraction of the intervals has the correct CN? This score is the percentage of intervals, weighted by length, that have the same CN in *S* and *T*. Unlike criteria 1-4, which are binary, this criterion measures the extent of correctness of a solution, and thus is more sensitive and accounts also for partially-correct solutions. We call it the *CN score*.

#### Base scenario

10,000 karyotypes were generated for the base scenario, and the algorithm was applied with bridge support weight *α* = 0.1. The performance is summarized in Figure.

To assess the distribution of each success rate criterion, the karyotypes were divided into 100 batches of 100 karyotypes each. Mean scores were captured for each batch and the variation of the mean was computed.

The algorithm correctly identified between 55% and 73% of the karyotypes in each batch, with an average of 62%. For an additional 13% of the cases, the solution had an equal CN profile as the correct solution, reaching a total of 75%. An average of 82% of all karyotypes resulted a solution with a score equal or better than the correct one. When disregarding missing bridges, the algorithm correctly identified an average 84% of karyotypes. The mean CN score of all the 10,000 simulations was 0.97 with a small standard deviation of 0.009.

#### The effect of separate parameters

The effect of separate parameters was tested by simulations in which one parameter was altered, while keeping the other parameters at their value in the base scenario. 100 simulated karyotypes were generated for each value and the percentage of solutions falling into the categories of correct, ECN, EBS and EOB was evaluated.

#### Bridge support weight

We first tested the effect of *α*, the relative weight assigned the bridges, on the performance, for 0 ≤ *α* ≤ 2. There is a noticeable improvement when *α* > 0, and little effect for the range of 0 < *α* ≤ 0.1. For larger values of *α* there is a small but noticeable negative effect. (Table S1).

#### Noise in copy number measurements

We tested the algorithm for different levels of CN noise *ε* under the base scenario. The results are shown in Figure 6. As expected, a higher level of noise makes it harder for the algorithm to find the correct solution. For *ϵ* ≤ 0.3 the performance of the algorithm is quite good, and for *ϵ* ≥ 0.4 the results begin to deteriorate. As expected, at high noise levels the majority of the solutions have better score than the true one.

#### The number of operations

We tested the algorithm on karyotypes that underwent 1 ≤ *N* ≤ 30 structural and numerical operations, under the base scenario. The results are shown in Figure 0. As expected, more operations make the problem harder and the success rates decrease. For example, the fraction of perfectly solved cases drops from 88% with one operation to less than 10% with 30 operations. The CN score drops more slowly, as CN of long fragments can still be reasonably inferred even if their order is incorrect.

#### Other parameters

When testing the effect of other parameters, the results met our expectations – karyotypes with less chromosomes (Figure S3) or a single copy of each chromosome instead of diploid (Figure S4) yielded better results. Results were also better when the probability of missing a bridge was lower (Figure S5).

We also looked at cancer heterogeneity situations. Different cells of the same tumor can have different karyotypes, having taken different evolutionary paths ^41–45^. Most cancer data today is still based on DNA from numerous cells, providing measurements from a mixture of genomes. Can the karyotype be reconstructed out of the heterogeneous mixture? When simulating data mixture of normal and a cancer karyotype results only dropped mildly with the relative abundance of normal data (Figure S8). However, when mixing two distinct cancer karyotypes, performance dropped rapidly with the heterogeneity (Figure S6).

Finally, we simulated karyotypes by selecting operations with frequencies as reported in ^46^ rounded to multiples of 10%. There was little difference in the success rates between the uniform distribution and the uneven one (Figure S7).

### Real tumor analysis

We applied the algorithm on data extracted from real samples. Malhotra et al. ^46^ examined whole genome sequencing data of 64 different tumor samples, and reported for each sample a CN profile and a set of bridges with their support. We first filtered from the data very small segments and the corresponding breakpoints (see Supplement). Often the set of normal chromosomes that are involved in rearrangements and CN changes in a tumor can be partitioned into independent groups of chromosomes (i.e., no two segments in different groups are connected by a bridge). In our graph representation, each such group is a connected component, which can be analyzed separately by the algorithm. The 64 tumor samples in ^46^ constituted together 570 such components, and each was analyzed separately.

#### Noise estimation

We first wanted to assess the noise level in the actual data affecting the reported CN values. Since CN in noiseless data should be integer, we estimated the noise *d_i_* for the reported CN *c_i_* as *c_i_* – [*c_i_*], where [*x*] is the nearest integer value to *x*. The CN data include 22,321 CN segments. A scatter plot of the standard deviation of the noise level vs. the number of bridges in each component can be seen in Figure 8. As expected, the mean noise level across the data was 0, showing that the noise is unbiased towards neither negative nor positive values. The standard deviation was 0.28, a value that we used as our default simulation scenario. Note that this estimate is a lower bound, since some measured CN values may actually differ from the real ones by more than 0.5.

In addition to CNs, the data include bridges and for each bridge an integer value, its support. The expected average support can be derived from the read depth and the insert size (see Supplement) and was found to be 10.7. The observed mean support score across all the data was 10.8. Figure 9 shows the distribution of the support scores across the data. The distribution closely resembles an exponential distribution with *λ* = 0.1866. For that reason, that was the value used in our simulation model (see supplement for more details).

#### The GBM10 sample

We analyzed in detail three components of bridge graphs obtained from real data. Table S2 shows information about them. Each has undergone 7-8 rearrangements, involving 1-4 chromosomes. For each component, the ILP algorithm outputs a directed weighted graph with a weight function that minimizes the distance and that can be broken into a set of paths *P* = {*p*_1_, … *p_n_*}, starting and ending at a telomere nodes, and alternating interval and non-interval edges. Another script translates the solution of the ILP solver to a dot language representation ^47^ that can then be visualized using a graph visualization tool such as GraphViz ^48^.

Figure 10A shows the graph corresponding to the component of chromosomes 4 and X in tumor sample GBM 10 (Glioblastoma multiforme). The resulting karyotype produced by our algorithm for this example is shown in Figure 10B. This graph can be broken into four different paths, representing both copies of the rearranged chromosomes 4 and X (Figure 10C).

The other two examples are described in the Supplement.

## Conclusions

In this work, the problem of inferring a tumor karyotypes from short paired end read data was investigated. A novel algorithm based on graph theory and ILP was introduced to solve the problem, and simulations were performed in order to evaluate the utility of such an approach. Some examples of analysis of real data were also presented.

To accurately estimate the correctness and robustness of the algorithm, validation against a data set of verified karyotypes is needed. However, a comprehensive set of sequenced tumor samples with CN profiles and paired-end reads data, matched with the corresponding true karyotypes, is currently not available. Data sets that currently exist either do not include a fully reconstructed karyotype, or include karyotypes of a very low resolution ^8^. We therefore used simulations to test and measure the success of our algorithm in a spectrum of scenarios, as well as to point out potential pitfalls.

The analysis of simulated data suggests that the most meaningful factors affecting the accuracy of solutions produced by our method are the noise and completeness levels of the data. We tested the algorithm in a scenario, designed to mimic parameter values observed in real data. Under these conditions, the algorithm correctly inferred 69% of the karyotypes. However, the success rate increased to 79% when considering solutions that are correct relative to the noisy input, and when accounting for unreported bridges, 87% of the tested cases were correct (Figure 5).

**Figure 5:**
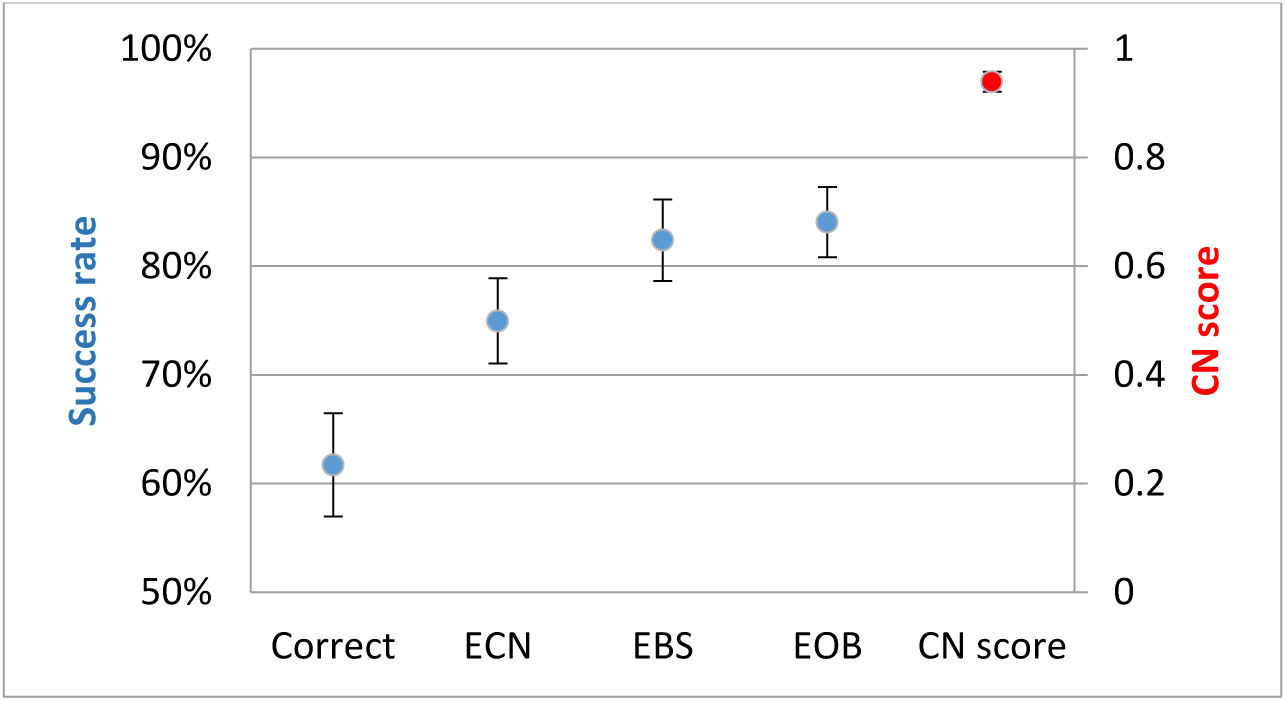
Distribution of the success rate over 100 independant simulations of the base scenario. Error bars are ± the standard deviation.

Furthermore, in scenarios where there is almost no noise, or when no bridges are unreported, the results are much better: accuracy was 90% and 100%, respectively (Figure 6, Figure S5). This strongly suggests that our method is limited mostly by the completeness and accuracy of the measured data. It suggests that more accurate sequencing technologies are needed in order to increase the chance to solve the karyotype reconstruction problem correctly.

**Figure 6:**
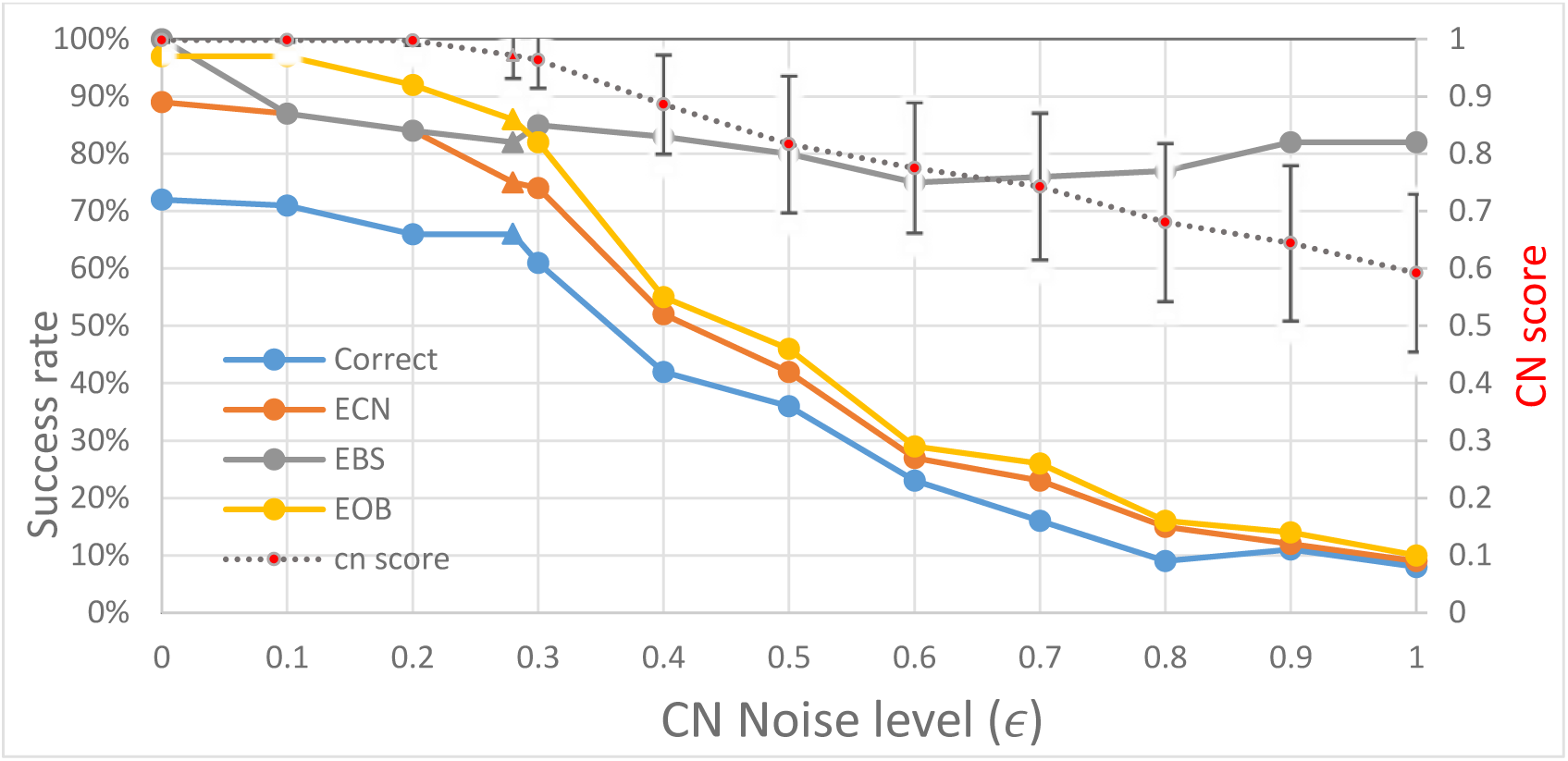
Performance of the algorithm as a function of noise level. For the CN score, the bars represent ±0.5 std. Data points for the default value of ϵ = 0.28 are marked with a triangle.

Our method was relatively robust when applied on data taken from tumor cells contaminated by healthy tissue (Figure S8). A sample that includes reads taken from a mixture of different tumor cells poses a bigger challenge, and the resulting karyotype is incorrect more often than it is correct (Figure S6).

Depending on one’s perspective, the results can be viewed as good or bad news. On the one hand, full, perfect reconstruction is not attained in over 30% of the cases. On the other hand, even in those imperfect cases, most of the reconstruction details are correct, as quantified by the other, less stringent, measurement criteria (Figure 5). Biological research has a great tradition of building up from incomplete data, the most obvious example being the human genome whose yet-incomplete versions have kept evolving for the past fifteen years. It may be the case that the imperfect reconstruction of cancer karyotypes can still produce valuable conclusions and findings.

### Limitations

Using simulations allows us to gain better understanding of the capabilities and limitations of our algorithm, but it requires us to make assumptions about the mechanisms driving genomic rearrangements in tumor cells and about the statistical properties of the read data. Both types of assumptions limit the generality of conclusions we can draw.

Firstly, our model defines a limited set of possible rearrangements (deletion, duplication, inversion and chromosomal translocation) and assumes that they occur with equal probabilities. Furthermore, our simulation of rearrangement events (except translocations) limits the genomic range they can span (see section 0) and assumes that events are equally likely to occur in any position on the genome. While these assumptions are very far from the real process of mutating cancer cells, they do provide a mechanism that can generate any rearranged karyotype. Our method proved robust when adjusting the frequency of each type of rearrangement to that observed in the data obtained from ^46^ (Figure S7), but other possible rearrangement mechanisms and their effect on the performance of the algorithm were not explored.

A second problem arises when attempting to create very complicated karyotypes using a large number of rearrangements. Stephens et al ^49^ suggested that in some cases a single catastrophic event called *chromothripsis* occurs, in which a section of the chromosome is shattered into a large number of small fragments and then reassembled, creating a karyotype that is much more complex ^49^. While all possible karyotypes can be generated using our model, very complex ones are unlikely. Note that once a deletion operation has been performed, the deleted segment cannot reappear and will therefore be absent from the final karyotype. When performing a large number of rearrangements on a chromosome, deletions will occur and sometime remove segments that were rearranged by a previous operation, essentially reducing the complexity of the resulting final karyotype. We tested our method on karyotypes that have undergone a maximum of 30 operations (Figure 0), but a modified simulation model needs to be used in order to generate more complex karyotypes. Currently our results reflect more faithfully the ability of the algorithm on relatively simple karyotypes, which constituted the majority in the real data that we used.

**Figure 0:**
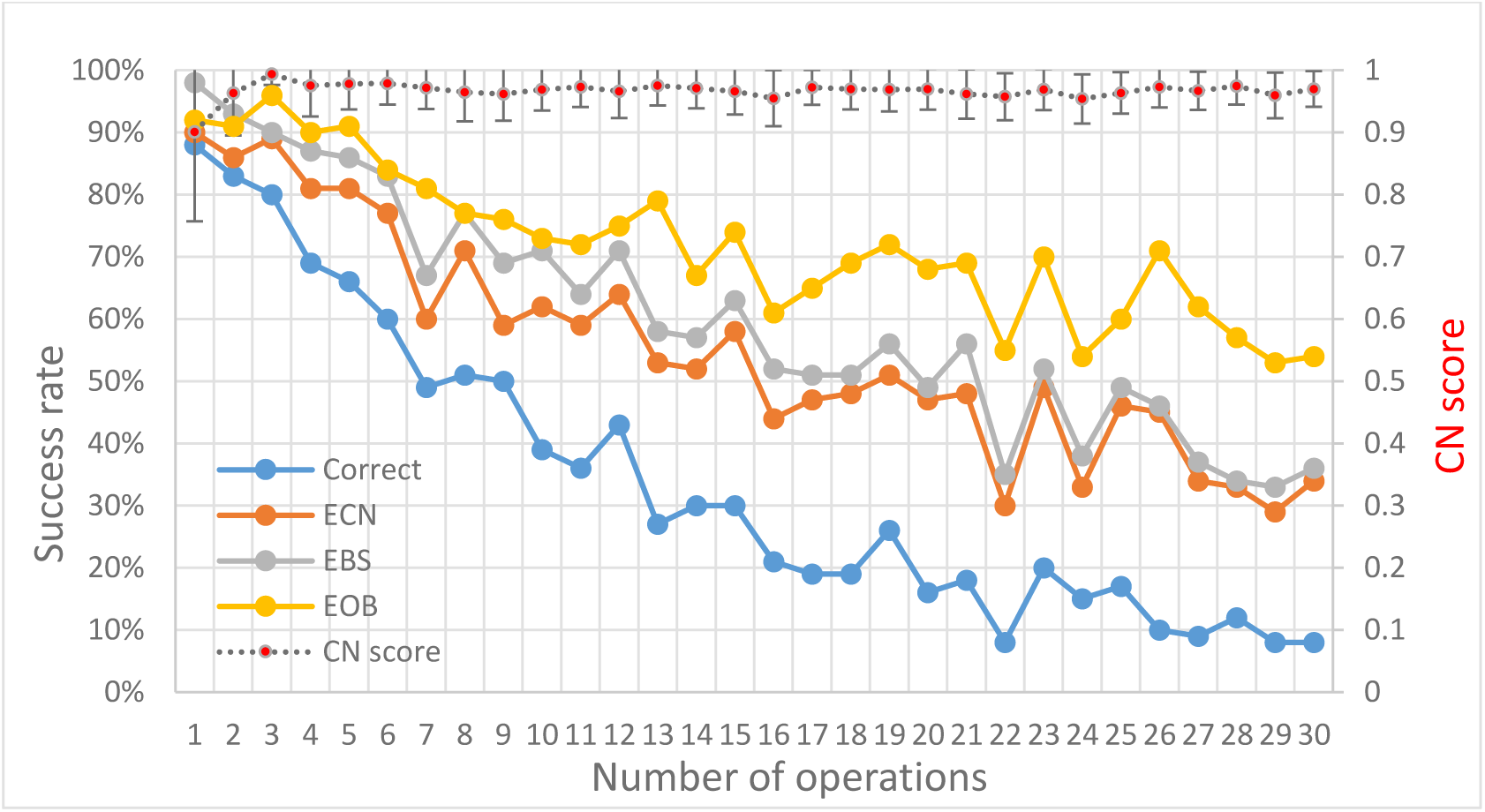
The effect of the number of operations. Success rates and CN scores. Error bars represent ±0.5 std.

**Figure 8:**
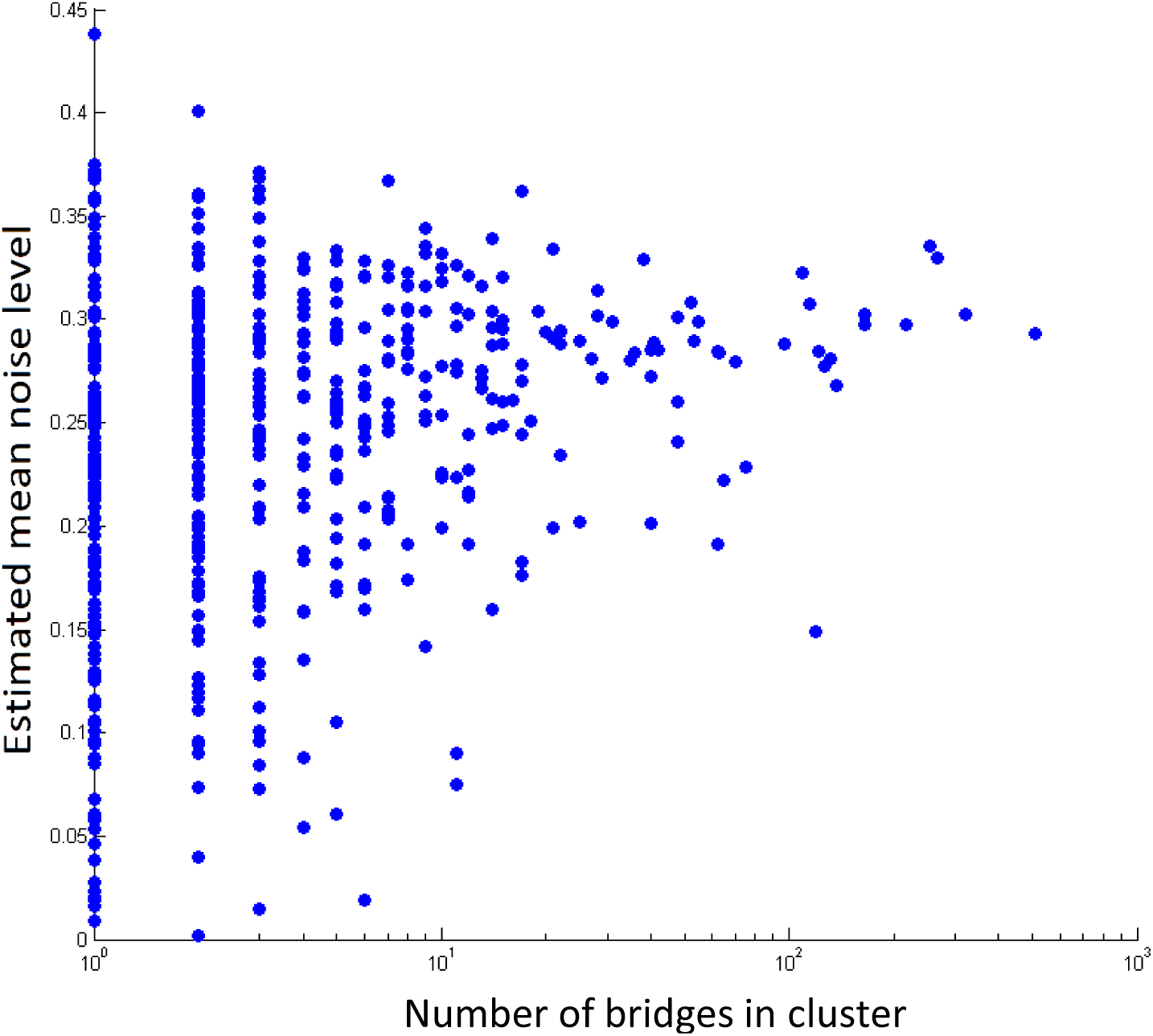
Estimated noise level in real cancer samples. The plot shows for each of the 670 components in the tumor samples in ^46^, the number of bridges and an estimate of the noise level calculated as standard deviation of the distances of the CN in the sample from the closest integer value.

**Figure 9:**
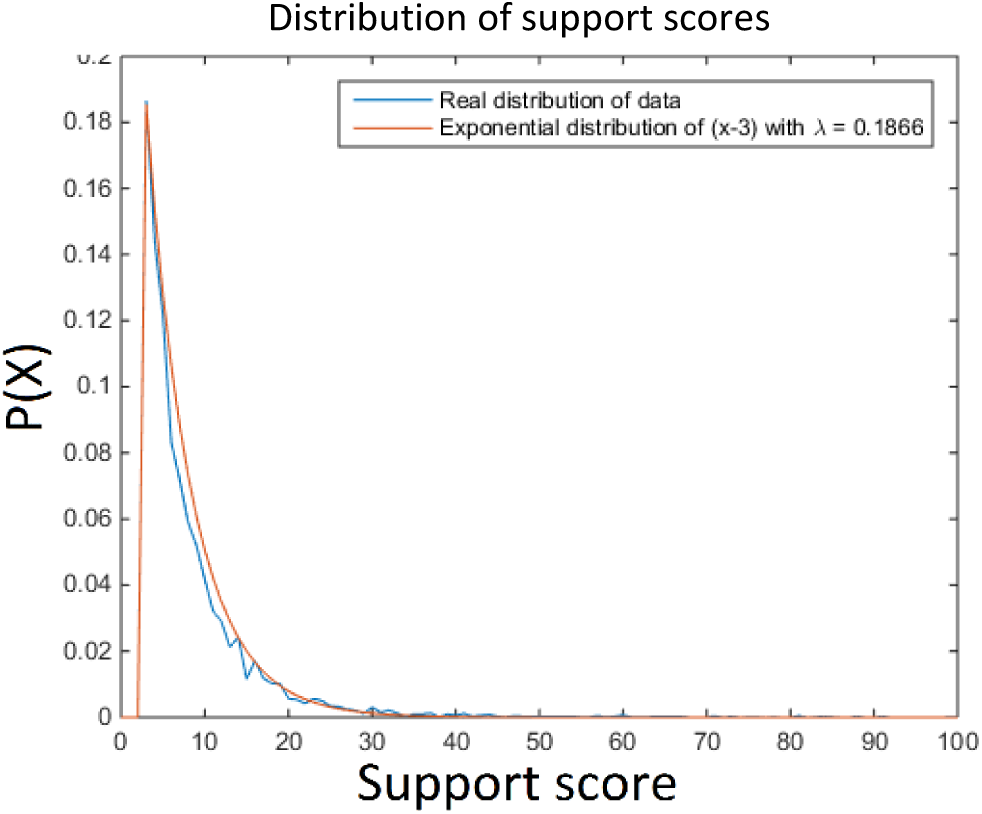
The distribution of the support score across the data plotted against an exponential distribution with λ = 0.1866. In both distributions values below 3 are ignored.

**Figure 10:**
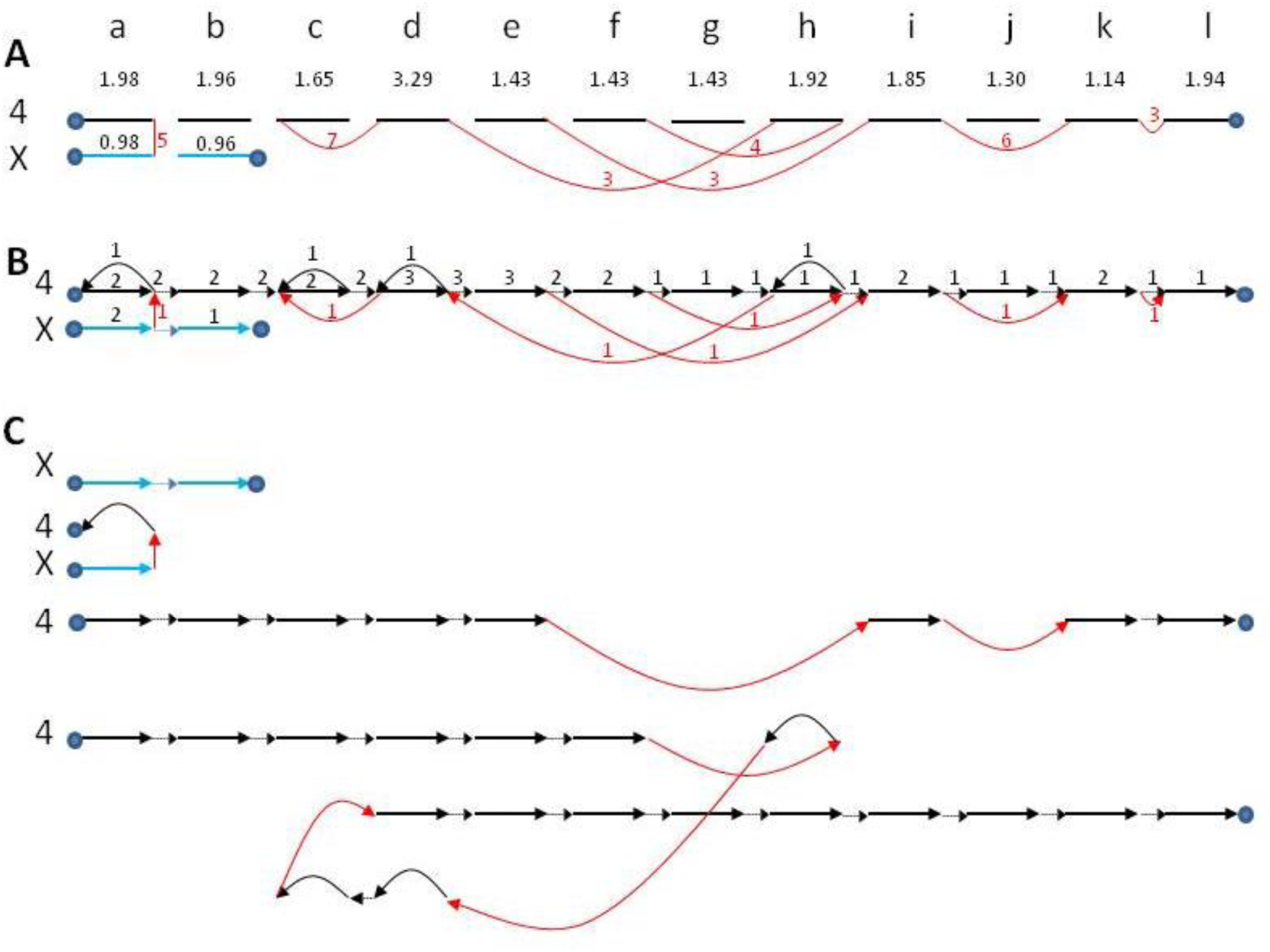
Results on sample GBM 10. The chromosomes were divided into segments according to the breakpoints inferred from the paired ends reads data and were named a-l. Segment sizes are not shown to scale. We mark interval, reference and bridge edges by black, dotted and red arcs respectively. The number next to a red edge (bridge) is the number of observed supporting reads for that bridge. In all subfigures the same intervals (here: a through l for Chr. 4 and a, b for Chr. X) are aligned. The numbers in the second line are observed coverage values. (A) Bridge graph for chromosomes X and 4. The bridge bteween segments k and l is a result of breakpoint filtering (see Supplement). (B) Solution suggested by our algorithm. For this sample the average distance of the resulting karyotype from the data, weighted by segment length, is 0.28. Note that segments a, c, d, and h have edges in both directions suggesting the solution includes traversal of these segments in both directions. (C) The different paths comprising the solution, representing the rearranged karyotype of chromosomes 4 and X.

A third type of limitation is due to the noise model assumptions. While we tried to borrow values of noise as estimated from the real data (see Real tumor analysis), there are other parameters that affect the noise and thus the quality of the analysis, including incorrectly mapped reads due to sequencing errors, non-uniquely mappable reads, insert length variance, breakpoints that fall within a read (and not in the gap), non-uniform read coverage, etc. These are all left to future work.

One of the limitations of our algorithm is its inability to “predict” bridges that were not observed in the data. The algorithm looks for a path on the graph corresponding to a karyotype that best fits the observed CN profile, yet it overlooks potential paths that can be constructed by bridging two unconnected interval edges – essentially predicting a bridge. This implies that data produced using sensitive methods, even with higher rates of false positives, might be preferable over data with false negatives.

### Future directions

One important aspect of the technology in detecting bridges is the insert size. A bridge will usually be detected only when the two reads of a PER are on the two different sides of it (see supplement). Therefore, the larger the read length and insert - the higher the bridge coverage. This implies that sequencing techniques with longer inserts can dramatically change the performance of the algorithm. Several such techniques are forthcoming, and some methods for detecting structural variations were already developed for them ^28^ ^29 22^. Note however that very short rearrangements that span less base pairs than the length of the read may be missed altogether.

A possible extension to our method can be the addition of weights to the reference edges. Recall that reference edges represent a connection between two segments that is expected according to the reference genome. Unlike interval edges or bridge edges, reference edges are weightless in our model. One metric that can be used to establish a confidence score for a reference edge is the number of PERs whose ends span the two segments bordering the reference connection.

## Declarations

### Abbreviations

ILP: Integer Linear Programming; CN: Copy Number; CNV: Copy Number Variations; PER: Paired-End Reads; ECN: Equal Copy Number; EBS: Equal or Better Score; EOB: Equal for Observed Bridges.

### Ethics approval and consent to participate

Not applicable.

### Consent for publication

Not applicable.

### Availability of data and material

All Data analyzed in this research was reported by Malhorta et al. in ^46^ and is available for download as supplemental material in http://genome.cshlp.org/content/23/5/762/suppl/DC1

### Authors’ contributions

RE and RS designed the study. RE prepared the data, developed the tools used to simulate and analyze data, and produced the results. RE and RS analyzed the results. RE and RS wrote the manuscript. Both authors read and approved the final manuscript.

## Acknowledgments

We would like to thank Nir Atias for help with the IBM CPLEX package and Roy Kasher for help with software design and optimization.

